# Thermophysiology or resource availability: What shapes the post-flooding abundance of lizard species across artificial islands in a Neotropical biome?

**DOI:** 10.1101/2025.09.12.675825

**Authors:** Rogério B. Miranda, Gideon G. Deme, Reuber A. Brandão

## Abstract

Global warming has led to increased flooding events, causing significant habitat fragmentation and affecting species distribution in tropical ecosystems. This study investigates the postflooding abundance of lizard species across artificially created islands in the Cerrado biome, focusing on the roles of thermophysiological traits and resource availability. We conducted a comprehensive survey, recording 560 individual lizards representing 13 species from eight families, and used generalized additive models to assess the impact of mean field-active body temperature and net primary productivity on lizard abundance. Our results indicate that a rise in mean field-active body temperature significantly correlates with reduced lizard abundance, supporting the physiological tolerance hypothesis. This finding points out the critical importance of thermoregulation for ectotherm survival in the face of climate change. Conversely, resource availability, measured through net primary productivity, showed no significant effect on lizard abundance. These findings bring to light the vulnerability of lizard populations to thermal stress and emphasize the necessity for conservation strategies that prioritize the creation of thermal refuges and improve habitat connectivity to mitigate the impacts of rising temperatures. By integrating species traits into macroecological analyses, our study offers valuable insights for biodiversity conservation in fragmented landscapes. Future research should explore diverse ecosystems to validate these findings, thereby informing effective conservation planning and ensuring the resilience of ectotherm populations in altered environments.

## Introduction

Global warming has significantly increased the risk and severity of natural disasters, like flooding, which has become more frequent due to the intensity of extreme precipitation events (Tabari, 2020; IPCC, 2021). The impact of flooding on the spatiotemporal distribution and redistribution of species has been suggested to largely depend on the severity and duration of the flood, and the ecosystem involved (Ozer et al., 2003). Thus, the complex and multifaceted impact of floods on the ecological landscape can directly or indirectly influence terrestrial biodiversity (Mahé et al., 2013). Although flooding can lead to habitat destruction and fragmentation, we have evidence suggesting that flooding also triggers breeding, migrations, and dispersal in some species (Zhang et al., 2021). Still, our understanding of the spatiotemporal distribution and redistribution of species across habitats created as a result of flooding is limited, especially in Neotropical ecosystems.

Habitat fragmentation, a by-product of flooding, represents one of the most relevant causes of biodiversity loss (Díaz et al., 2022). This is because fragmentation, leading to the creation of “new” habitats, can alter the structure and composition of ecosystems, reduce the availability of resources, and impact the survival strategies of species (Ebrahimi et al., 2023). Thus, habitat reorganization can influence underlying changes in patterns of diversity and distribution of organisms, which may increase the extinction of certain species (Diamond, 1989) within and among biomes across local, regional, and global scales (Hansen et al., 2010). For example, species’ survival rates were found to be higher with a lower number of patches found across larger areas (Niebuhr et al., 2015). Perhaps, the potential explanation for this ecological phenomenon is the species–energy theory, where resource availability drives the spatial population dynamics of species (Wright, 1983), with a secondary role for thermophysiology in determining species’ spatial distribution patterns across habitat patches along gradients (Currie et al., 2004). Previous studies have tested the species–energy theory to explain the spatial distribution pattern of various species across habitat patches (Srivastava and Lawton, 1998; Currie et al., 2004; Evans et al., 2005; Buckley and Jetz, 2010; Vagle and McCain, 2020). However, these studies have found mixed support for different groups of species across gradients. For instance, the extinction rates of the British avifauna were found to be slightly lower in high-energy areas with more abundant species, demonstrating little support for the species–energy theory (Evans et al., 2005). Meanwhile, there was no such relationship between resource availability and the spatial variations of American avifauna or butterflies (Currie et al., 2004). Rather, the spatial variations of American avifauna or butterflies were found to vary according to the tolerances of species to environmental conditions (Currie et al., 2004), suggesting the ‘physiological tolerance hypothesis’ in explaining the species-abundance patterns (Currie et al., 2004) across habitat patches along gradients. Given the important implications of species–area in conservation planning and ecosystem management for altered landscapes, including the design of areas to enhance habitat connectivity and population viability (Vagle and McCain, 2020), understanding how species interact with the environment in fragmented habitats becomes a priority. This is even more important for ectotherms like lizards, whose distributions are known to be sharply constrained by prevailing environmental conditions (Gainsbury and Colli, 2019).

Lizards face heightened extinction risks due to rising temperatures (Sinervo et al., 2010; Pontes-da-Silva et al., 2018). As ectotherms, lizards’ body temperature is a key factor in all physiological and behavioral processes (Huey, 1982). Recent studies suggest that tropical lizards may already be experiencing temperatures that surpass their physiological limits, thereby elevating extinction threats (Huey et al., 2009; Diele-Viegas et al., 2018). The risk of extirpation or extinction arises from the restriction in activity time for performing essential functions for survival (Buckley and Jetz, 2010; Sinervo et al., 2010; Diele-Viegas et al., 2018). In order to survive these new environmental conditions, lizards may adjust by behavioral and physiological plasticity or adaptation (Sinervo et al., 2010). Behavioral thermoregulation, such as basking and altering habitat preferences, allows lizards to optimize performance (Li et al., 2017). However, due to phylogenetic constraints, lizards’ active body temperature (T_b_) cannot evolve at the same pace as climate change (Sinervo et al., 2010). Perhaps, because of plasticity or adaptation (Sinervo et al., 2010), both resource availability and behavioral thermoregulation are able to relate to the variation of lizards’ total abundance across habitats. Thus, there might be a cascading effect of local resources partitioning in fragmented habitats occupied by species (MacArthur, 1960), and environmental temperatures, which strongly constrain lizards’ physiology (Huey, 1982), on the geographic distributions at local scales (Huey et al., 2012).

Here, we used the tropical lizards, known for maintaining cool temperatures in low-latitude regions where shaded areas play a crucial role in buffering climate warming effects (Kearney et al., 2009) as a priority, to assess the specific determinants (thermophylogical trait vs resource availability) in the variation of species abundance across artificially created islands post flooding in the Cerrado biome. In this study, we aim to understand how thermophysiological traits of lizard species in the Cerrado biome, together with resource availability, influence species’ total abundance in fragmented habitats. Unsuitable temperatures and habitat fragmentation can increase organismal vulnerability (Huey et al., 2012), particularly among lizard groups with different life-history traits occurring around the Serra da Mesa hydroelectric sites (Miranda et al., 2024). Specifically, we aim to investigate the species–resource availability hypothesis (Wright, 1983) vs the physiological tolerance hypothesis (Currie et al., 2004), to understand the spatial population dynamics of lizard assemblages in the fragmented habitats. On the species–resource availability hypothesis, with the evidence that lizards have much lower energy requirements compared to endotherms (Pough, 1980), we expect likely little or no relationship between local net primary productivity (NPP) and the number of individuals across lizard communities, with lizards’ total abundance expected to be at most weakly related to habitat productivity. On the other hand, on the physiological tolerance hypothesis, we expect thermophysiological traits (i.e., body temperature) to influence the spatial demography and ultimately the abundance of tropical lizard species. This is because thermoregulation can buffer the effects of prevailing environmental conditions (Huey et al., 2003), assuming that behavioral thermoregulation remains effective when the energetic demands are equivalent among species within a community (Buckley et al., 2008).

## Methods

### Study area

In the Cerrado biome, the Serra da Mesa dam in Goiás, Central Brazil (48º 20’W; 13º 51’S), created a 178 km^2^ lake by damming the Tocantins River. From 1996 to 1999, this flooding isolated 280 hilltops as islands, 80% of which are under 3 ha, with some exceeding 1000 ha. Eight hills were chosen for pre-flood monitoring, and after flooding, five islands (I-34, I-35, I-37, I-38, IX; Fig.1) remained, ranging from 3 to 35 ha. Selection was based on vegetation type, disturbance levels, and accessibility. Island IX later reconnected to the mainland due to fluctuating water levels.

**Figure 1.**
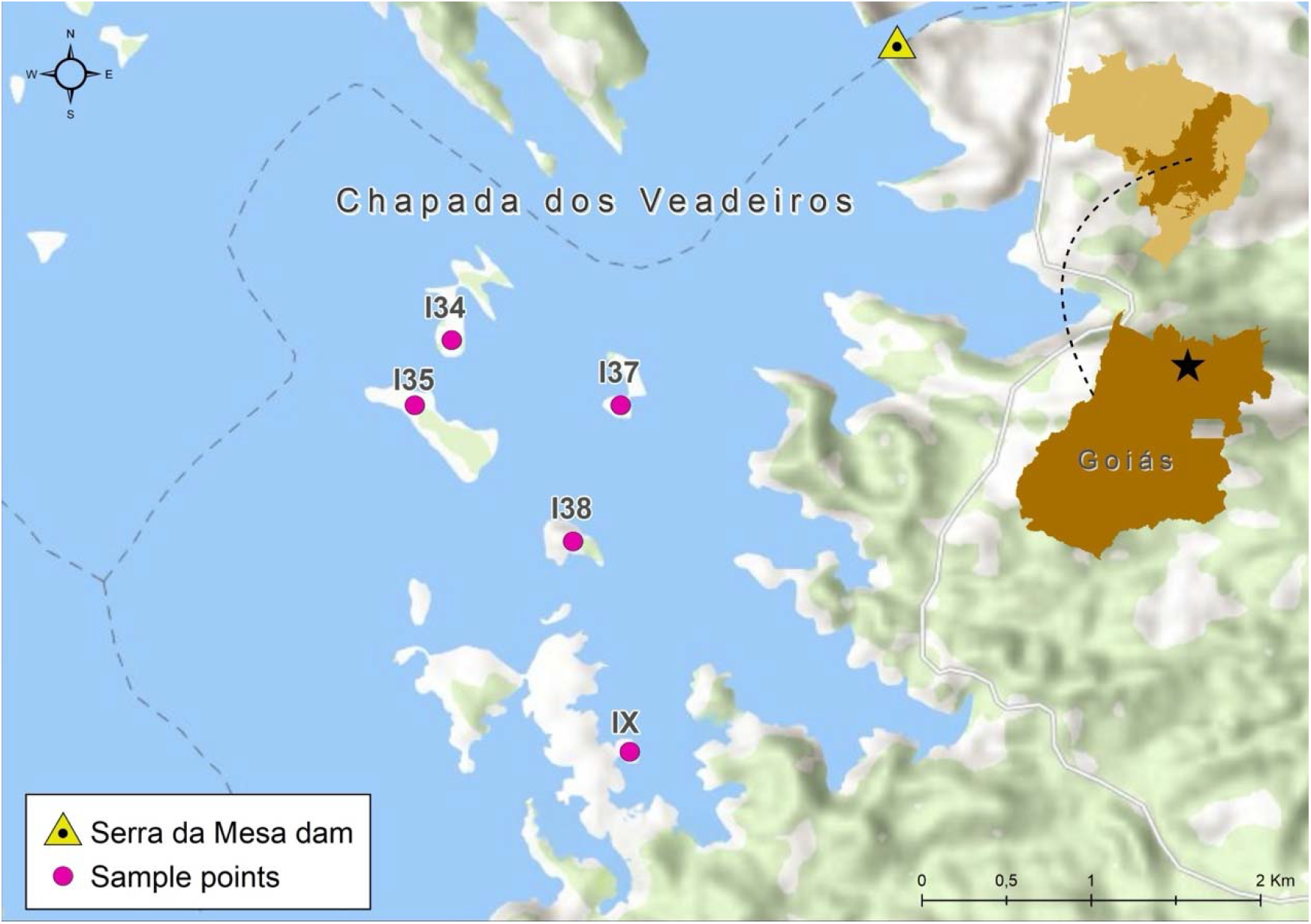
Map of the Serra da Mesa dam and lake in Goiás State, Brazil, showing the monitoring sites. Insets show a map of Brazil (upper) with the Cerrado region shaded, and the state of Goiás (lower), which is centrally located in this region.

### Species survey and monitoring

After the flooding in 2001, rocky soils and limited sample areas required the use of fire square sampling. Ten 50×50m plots, enclosed with plastic fences, were cleared using controlled burns, which exposed lizard shelters. Since the sampling effects on species richness and composition were controlled using abundance-based rarefaction curves effects of sampling (Brandão 2002), we are confident that this methodology presents satisfactory efficiency. Moreover, this methodology was also successfully used in previous studies (e.g., Amorim 2017; Miranda et al., 2024) with satisfactory results. Searches continued until no lizards were found for two hours. Lizards were measured, weighed, euthanized with lidocaine, preserved, and deposited in Coleção Herpetológica da Universidade de Brasília (CHUNB). This method ensured no injuries and effective capture.

### Thermophysiological data

The thermophysiological parameter used in this study was the mean field-active body temperature (T_b_). We obtained T_b_ for all species from Dielle-Viegas et al. (2018), except for one, *Tropidurus montanus*, which had its T_b_ acquired from Kiefer et al. (2005). In ectotherms, all physiological and behavioral processes are tied to T_b_; therefore, its maintenance within an adequate range is critical for their survival (Huey, 1982; Dielle-Viegas et al., 2018). Moreover, other essential activities for lizards’ survival are reliant on T_b_, such as reproduction, digestion, and foraging (Huey, 1982).

### Resource availability across islands

We used the Raster package in R to extract available resources in terms of net primary productivity (NPP) for each of the artificially created islands occupied by lizards (Hijmans & Etten, 2012). In extracting NPP as a variable, we used the highest resolution within a 2.5 arc min resolution grid (1□ ×□1□km) from the variables extracted from the Climatologies at high resolution for the earth’s land surface areas (CHELSA; https://chelsa-climate.org/bioclim/v2.1 accessed on May 30, 2025), which allowed us to obtain the calculated total plant biomass minus the carbon lost to respiration measured in gC□m−2□year−1 from Earth’s land surface areas for each of the islands. By using NPP as the proxy to measure resource availability for species (Huston & Wolverton, 2011; Meiri et al., 2007), we are able to collect information on the spatiotemporal variability in available resources across islands.

### Statistical Analyzes

All data visualizations and analyses were performed using the R programming language version 4.3.0 (www.r-project.org). Since lizard species abundance recorded the number of individuals across the years of the study, we computed the presence-absence matrix to allow us to accurately analyze the spatiotemporal distribution of lizard patterns across islands. By summing up the computed presence-absence matrix of lizard species, were able to calculate the number of species found in each island, which we used as response variables in our models, to assess the influence of island creation due to flooding, thermophysiology, and the variability in resource availability on the spatiotemporal distribution of patterns lizards across the artificially created islands in our study location.

To specifically determine how the field-active body temperature (thermophysiology of lizards) and resource availability are going to differently influence the spatiotemporal variability in lizards’ true abundance across islands, we fitted the thermophysiology of lizards and resource availability in different Generalized Additive Models (GAMs), while having island types as a fixed factor in both models. We fitted these factors in different models because of the evidence demonstrating that the variability in environmental conditions, which strongly relates to available resources across habitats, can influence lizards’ most active time to scout for resources (Meiri et al., 2013, 2020). For the GAM models, we used the approach without the smooth term on our predictor variables in each model (islands and thermophysiology vs islands and resource availability model), to assess the linear and non-linear relationships between lizards’ true abundance and the predictor variables across isolated local gradients, such as the artificially created islands. In total, we ran two GAMs, with each having species abundance indexes as the response variable, and islands with thermophysiology or islands with resource availability model as predictor variables, while accounting for individual species variation as a random variable. In each model, we log-transformed our predictor variables to improve the residual normality and reduce heteroscedasticity of our data. We used the Akaike Information Criteria (AIC) to evaluate our fitted models for the best-fitted model, while using the ANOVA for the GAMs approach for our hypothesis testing (Hastie, 2017) to determine the specific factors influencing lizards’ species abundance across artificially created islands, post-flooding.

## Results

A total of 560 individual lizards were recorded, representing 13 species from eight families: Teiidae (3 species), Gymnophtalmidae (2), Scincidae (2), Tropiduridae (2), Dactyloidae (1), Iguanidae (1), Phyllodactylidae (1), and Polychrotidae (1) (Table S1). During the study period, we observed variations in the individual lizard species recorded across five different artificially created islands and their environs. The most encountered lizard species was the *Copeoglossum nigropunctatum*, with 168 individuals recorded for this species. In contrast, the least encountered lizard species were the *Anolis brasiliensis, Polychrus acutirostris*, and *Tupinambis quadrilineatus*, with one (1) individual recorded for each of these species.

We found that lizards’ true abundance was significantly influenced by the mean field-active body temperature (mean field-active body temperature: β = −16.642 ± 6.077, χ2 = 7.500, *p* = 0.006) of lizard species post-flooding, with the abundance of lizard species declining as the mean field-active body temperature of individual lizard species increases (Fig. 2). However, we did not find any evidence to suggest that the decline in the abundance of lizard species was either influenced by the artificially created island types or the net primary productivity of these artificially created islands (Fig. 3; Table 1).

**Table 1.** ANOVA results and parameter estimates of lizard species total abundance as a function of thermophysiology (lizards’ active body temperature) and resource availability predictors. (a) ANOVA table for a model with lizards’ active body temperature (T_b_), and (b) ANOVA table for a model with net primary productivity (NPP).

| Factor | d.f. | $\chi^2$ | $p$ |
| --- | --- | --- | --- |
| a. Model with $T_b$ | | | |
| Island type | 4 | 0.928 | 0.920 |
| $\log(T_b)$ | 1 | 7.500 | 0.006 |
| AIC | 12.891 | 81.796 |  |
| Residuals | -0.367 | 0.351 |  |
| b. Model with NPP |  |  |  |
| Island type | 4 | 1.456 | 0.834 |
| $\log(NPP)$ | 1 | 0.004 | 0.950 |
| AIC | 12.972 | 82.021 |  |
| Residuals | -0.246 | 0.413 |  |

Note: Individual lizard species abundance had significantly high non-linear relationships with the predictor variables in both models: for (a) the effective degrees of freedom (edf) estimated = 6.415, $\chi^2 = 520.2$ , $p < 0.0001$ ; for (b) the effective degrees of freedom (edf) estimated = 6.699, $\chi^2 = 320$ , $p < 0.0001$ . There was no significant deviation explained (d.f. = -0.121, deviance = -0.063, $p = 0.164$ ) between the models with thermophysiology and resource availability predictor variables, demonstrating that both models had a similar good fit.

**Figure 2.**
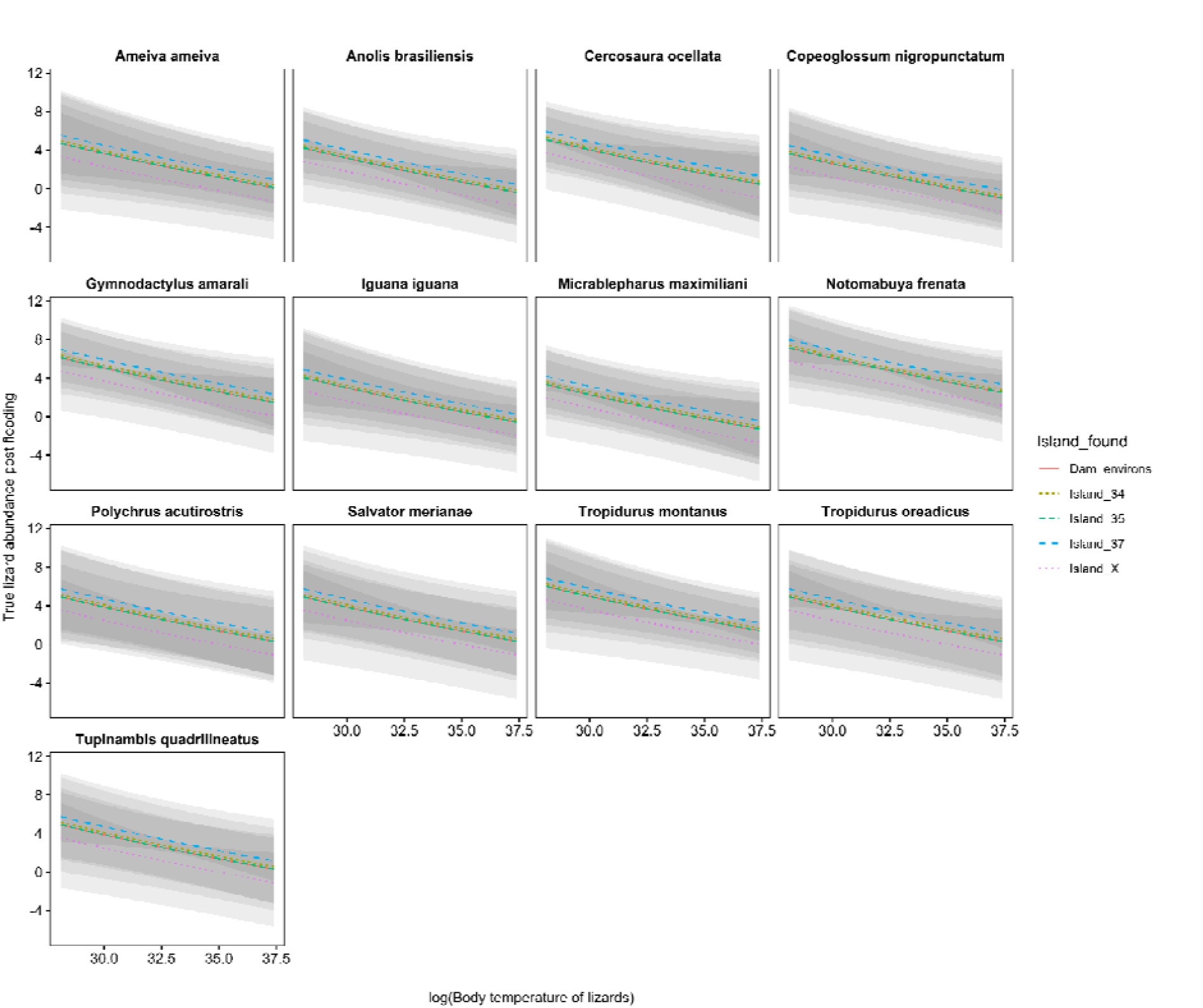
Results of our analysis for lizard species’ abundance across artificially created island with body temperature of lizards. The y-axis is represented as the magnitude and pattern of changes of lizards’ abundance, with a soothing effect of the body temperature of lizards and amily-level across island types. Grey-colored lines and ribbons represent the predicted values ± 1 SE of estimates of GAMs.

**Figure 3.**
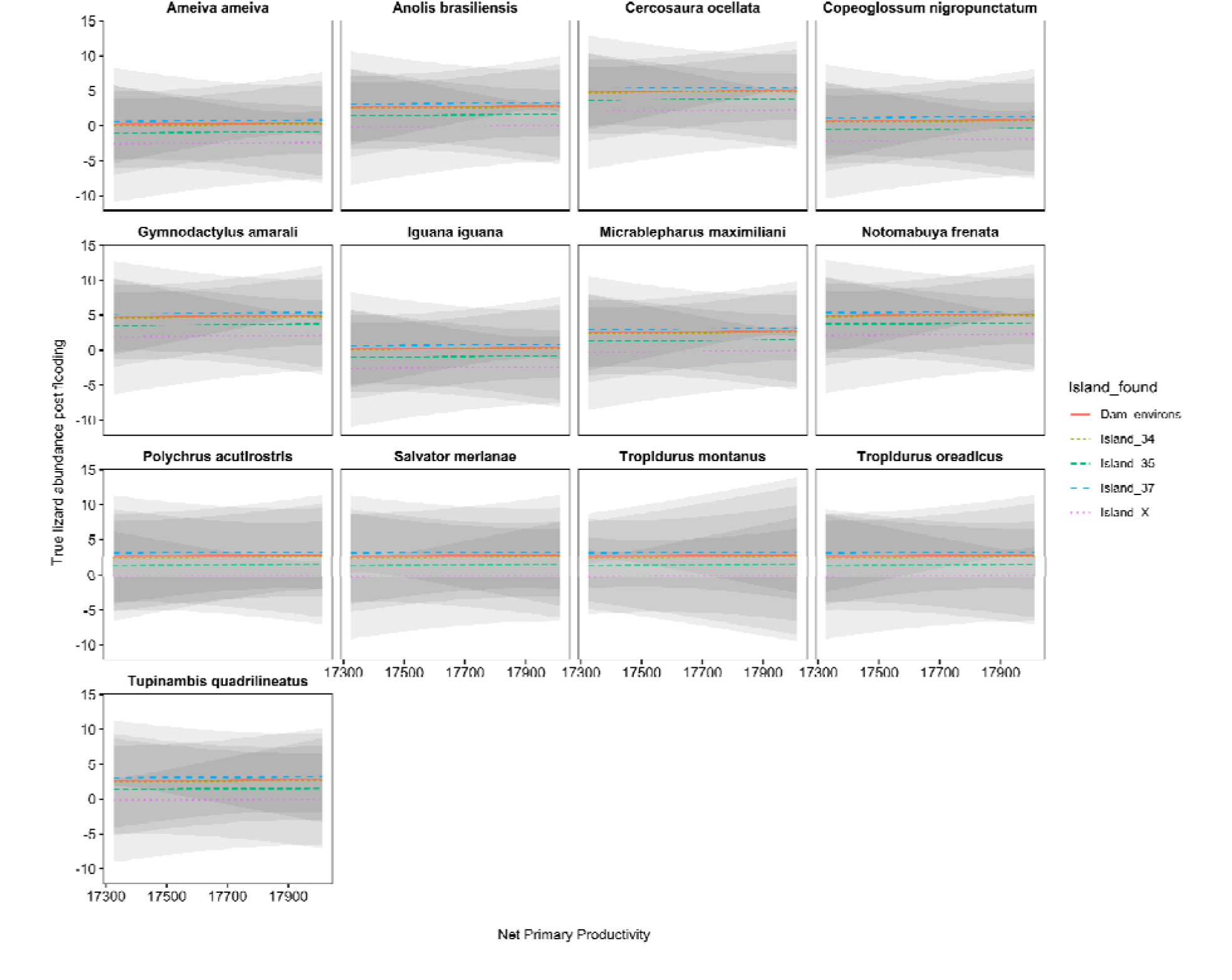
Results of our analysis for lizard species’ abundance across artificially created island with net primary productivity across different islands. The y-axis is represented as the magnitude and pattern of changes of lizards’ abundance, with a soothing effect of net primary productivity and family-level of lizards across island types. Grey-coloured lines and ribbons represent the predicted values ± 1 SE of estimates of GAMs.

Our model performance assessment and comparison indicated that the mean field-active body temperature of individual lizard species was probably the ecological factor influencing the abundance of lizard species in our study area, because our best-fit model was the model with island types and the mean field-active body temperature of individual lizards as predictor variables (AIC = 81.7958; Adjusted R-squared = 0.999; Table 2).

## Discussion

It has become important to link species traits in macroecological patterns’ analysis, as it offers a promising means to better identify the specific determinants of species community recomposition dynamics in post-anthropogenic change scenarios (Chapin et al., 2000; Cadott et al., 2011; Violle et al., 2014; Gaüzère et al., 2025). Our results demonstrate a significant decline in lizard abundance with increasing mean field-active body temperature, which suggests that thermoregulation was crucial in determining species distribution and abundance post-flooding. Importantly, the findings from our study offer valuable insights about the role of thermophysiological traits in shaping the abundance of tropical lizard species in fragmented habitats. Further, we find support for the physiological tolerance hypothesis, highlighting the constraints imposed by abiotic environmental conditions on lizard populations. Indeed, the physiological tolerance hypothesis supported in our study aligns with previous research emphasizing the importance of thermoregulation for lizards’ survival in a rapidly warming scenario (Huey et al., 2003; Buckley et al., 2008). The implication of our study shows that encompassing species traits (Cadott et al., 2011; Gaüzère et al., 2025), like habitat and climatic preferences of ectotherms (i.e., thermophysiology), can robustly indicate ectotherm species’ susceptibility or adaptability to anthropogenic climate change impacts.

By linking lizard species thermophysiological traits in our study, we aim to highlight how trait-based macroecological patterns, even when using sparse species long-term occurrence data, can provide a more sensitive depth for detecting anthropogenic change impact on biodiversity trends, especially among ectotherms with high environmental sensitivity (Sinervo et al., 2010; Diele-Viegas et al., 2018; Carmona et al., 2021). Indeed, our study found a negative correlation between body temperature and species abundance, which points out the vulnerability of lizards to rising temperatures. Our findings are in divergence from the species–energy theory but align with findings by Currie et al. (2004) that found physiological constraints as more influential than energy-driven dynamics in tropical lizard communities. Moreover, contrary to the species– energy theory, which states a relationship between resource availability and species abundance, our results indicate that resource availability, measured through net primary productivity, does not significantly impact lizard abundance. In a previous work, Miranda et al. (2024) speculate that species with lower energy demands were favored over the larger ones since only smaller lizard species prevailed on the islands. The current results also refute that possible explanation for the local extinction process.

In summary, our study confirms our prediction that thermophysiological traits are the main drivers in determining the abundance of tropical lizard species in fragmented habitats. While our study provides valuable insights, we find it important to acknowledge some limitations. The scope was confined to artificially created islands in the Cerrado biome, which may not fully represent other Neotropical ecosystems. Future research should expand to different habitat types and geographical regions to validate the generalizability of our findings.

## Implications for Conservation

Our findings present profound implications for conservation planning and ecosystem management. We highlight the sensitivity of lizard species to thermal conditions and the need for prioritization of habitat designs that enhance thermal refuges and connectivity, particularly in flood-prone areas. Conservation strategies should focus on mitigating temperature extremes and preserving shaded microhabitats to buffer climate warming effects (Kearney et al., 2009; Cadott et al., 2011). Additionally, by advancing our understanding of species–environment interactions, we can better inform conservation strategies to safeguard biodiversity in altered landscapes and mitigate the impacts of human activities.

## Acknowledgments

We thank Maria Eduarda Coelho for the help with the map. This work was funded by the Presidential Society of STEM Postdoctoral Fellows program at Case Western Reserve University to RBM, and Centro UnB-Cerrado to RAB and RBM. RAB also thanks CNPq (Conselho Nacional de Desenvolvimento Científico e Tecnológico) for his research fellowships and financial support (processes 306644/2023-7 and 306994/2023-2).

