## Supplemental Material for "Thermophysiology or resource availability: What shapes the post-flooding abundance of lizard species across artificial islands in a Neotropical biome?"

Supplementary information

Table S1. Species list in family alphabetical order.

| Species | Family |
| --- | --- |
| *Anolis brasiliensis* | Dactyloidae |
| *Cercosaura ocellata* | Gymnophtalmidae |
| *Micrablepharus maximiliani* | Gymnophtalmidae |
| *Iguana iguana* | Iguanidae |
| *Gymnodactylus amarali* | Phyllodactylidae |
| *Polychrus acutirostris* | Polychrotidae |
| *Copeoglossum nigropunctatum* | Scincidae |
| *Notomabuya frenata* | Scincidae |
| *Ameiva ameiva* | Teiidae |
| *Salvator merianae* | Teiidae |
| *Tupinambis quadrilineatus* | Teiidae |
| *Tropidurus montanus* | Tropiduridae |
| *Tropidurus oreadicus* | Tropiduridae |

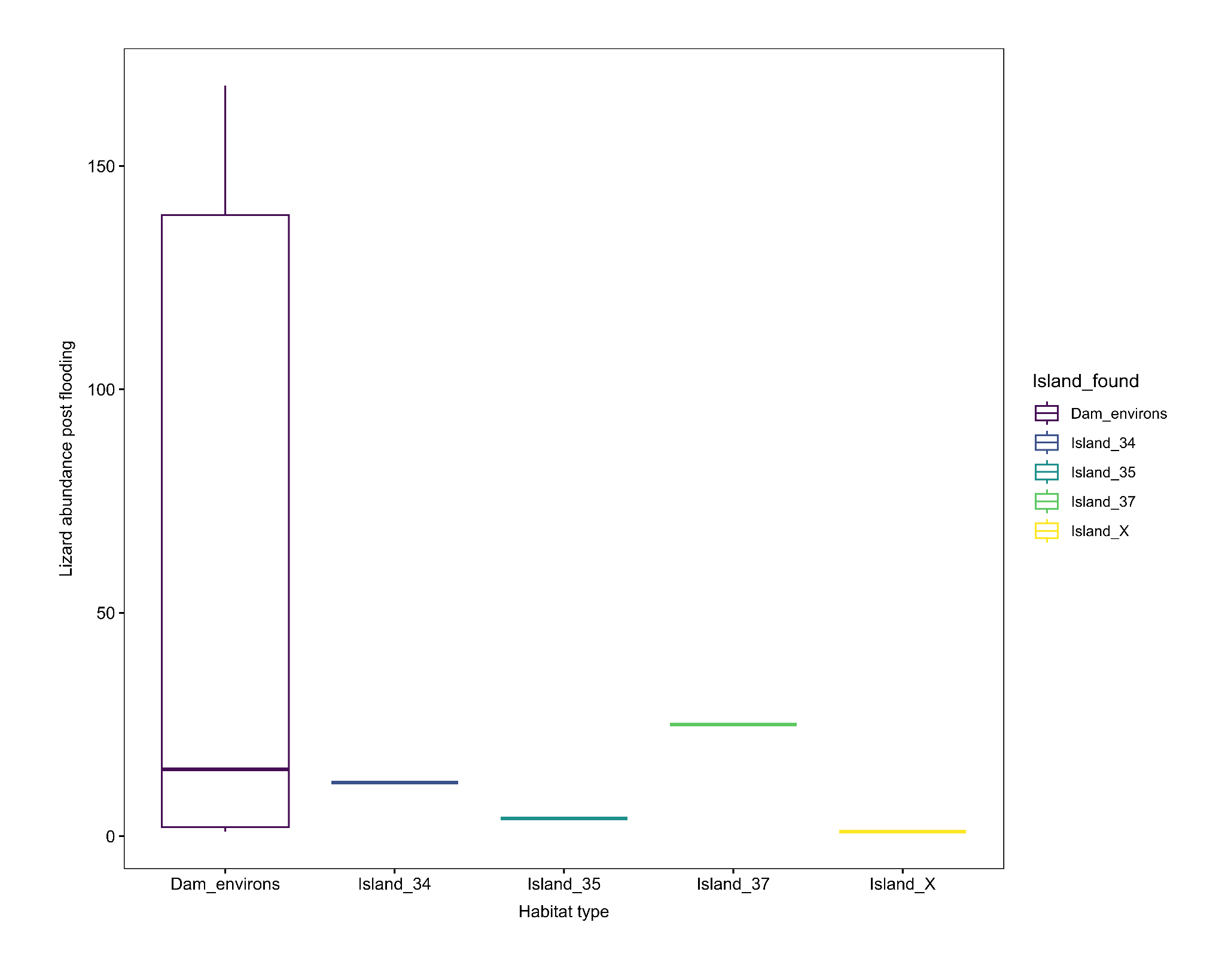

Fig. S1. Boxplot showing lizard species’ abundance across the different islands post-

flooding
